# Elucidating the dynamics of Integrin αIIbβ3 from native platelet membranes by cryo-EM with build and retrieve method

**DOI:** 10.1101/2024.11.27.625729

**Authors:** Xu Han, Zhemin Zhang, Chih-Chia Su, Meinan Lyu, Masaru Miyagi, Edward Yu, Marvin T. Nieman

## Abstract

Platelets fulfill their essential physiological roles sensing the extracellular environment through their membrane proteins. The native membrane environment provides essential regulatory cues that impact the protein structure and mechanism of action. Single-particle cryogenic electron microscopy (cryo-EM) has transformed structural biology by allowing high-resolution structures of membrane proteins to be solved from homogeneous samples. Our recent breakthroughs in data processing now make it feasible to obtain atomic-level-resolution protein structures from crude preparations in their native environments by integrating cryo-EM with the “Build-and-Retrieve” (BaR) data processing methodology. We applied this iterative bottom-up methodology on resting human platelet membranes for an in-depth systems biology approach to uncover how lipids, metal binding, post-translational modifications, and co-factor associations in the native environment regulate platelet function at the molecular level. Here, we report using cryo-EM followed by the BaR method to solve the unmodified integrin αIIbβ3 structure directly from resting human platelet membranes in its inactivated and intermediate states at 2.75Å and 2.67Å, respectively. Further, we also solved a novel dimer conformation of αIIbβ3 at 2.85Å formed by two intermediate-states of αIIbβ3. This may indicate a previously unknown self-regulatory mechanism of αIIbβ3 in its native environment.

In conclusion, our data show the power of using cryo-EM with the BaR method to determine three distinct structures including a novel dimer directly from natural sources. This approach allows us to identify unrecognized regulation mechanisms for proteins without artifacts due to purification processes. These data have the potential to enrich our understanding of platelet signaling circuitry.

**Key points:**

- We report the first structural analysis of platelet membrane proteins extracted directly from human platelet membranes.
- Our novel structural-omics approach allowed us to solve integrin αIIbβ3 structures in two distinct states from resting human platelets.
- Regulatory cues of integrin αIIbβ3 were preserved from the native resource and revealed on the models with atomic resolutions.
- This study opens the potential to build the platelet membrane protein atlas to understand platelet physiology.

## Introduction

Platelets are central players in primary hemostasis, thrombosis, inflammation, and vascular biology.^1,2^ Additionally, platelets have critical roles in diverse physiological and pathological processes ranging from inflammation and host defense to cancer, neurodegenerative, and cardiovascular diseases.^3–5^ Platelet membrane proteins serve as the first responders to environmental changes by relaying the extracellular information across the membrane to mediate the platelet response.^5–7^ While platelets patrol the integrity of the vascular system, membrane proteins sense and are regulated by their microenvironment. Establishing the interaction network of platelet membrane proteins will further our understanding of platelet responses.

Structural biology approaches like crystallography and cryo-electron microscopy (cryo-EM) have uncovered the functions of platelet membrane proteins by obtaining high-resolution structures of G-protein coupled receptors, adhesion molecules, and other surface receptors.^8–10^ They have also uncovered the binding sites and mechanisms of anti-platelet therapies. ^11,12^ However, to understand the impact of native environment, we need approaches beyond traditional structural biology. A combination of proteomics and high-resolution unmodified protein complex structures is essential to significantly advance our understanding of protein function *in situ*. Traditional proteomics offers valuable network insights but lacks high-resolution structural information. ^13,14^ Similarly, traditional structural approaches require highly purified homogenous protein samples, limiting their application in studying integrated systems within cells, tissues, or organs. ^15^ Moreover, the dynamics of membrane proteins pose further challenges to obtaining structural information without introducing stabilizing modifications in overexpression systems. ^16–18^ Finally, extensive purification of exogenously expressed proteins can lead to the loss of native regulatory cues and rare protein conformations.

Recent advancements in data processing now present the opportunity to bridge these gaps enabling us to gain unprecedented insights into the intricate workings of platelet membrane proteins in their native environment. These cutting-edge developments in the data processing algorithm make using cryo-EM to study whole-cell, tissue, or organ samples feasible.^19–23^ A novel iterative methodology, Build-and-Retrieve (BaR), was developed to process heterogeneous cryo-EM sample datasets (**Figure 1**).^21^ This methodology allows us to analyze crude samples by performing *in silico* purification during data processing from a large heterogeneous dataset. ^19–23^

**Figure 1.**
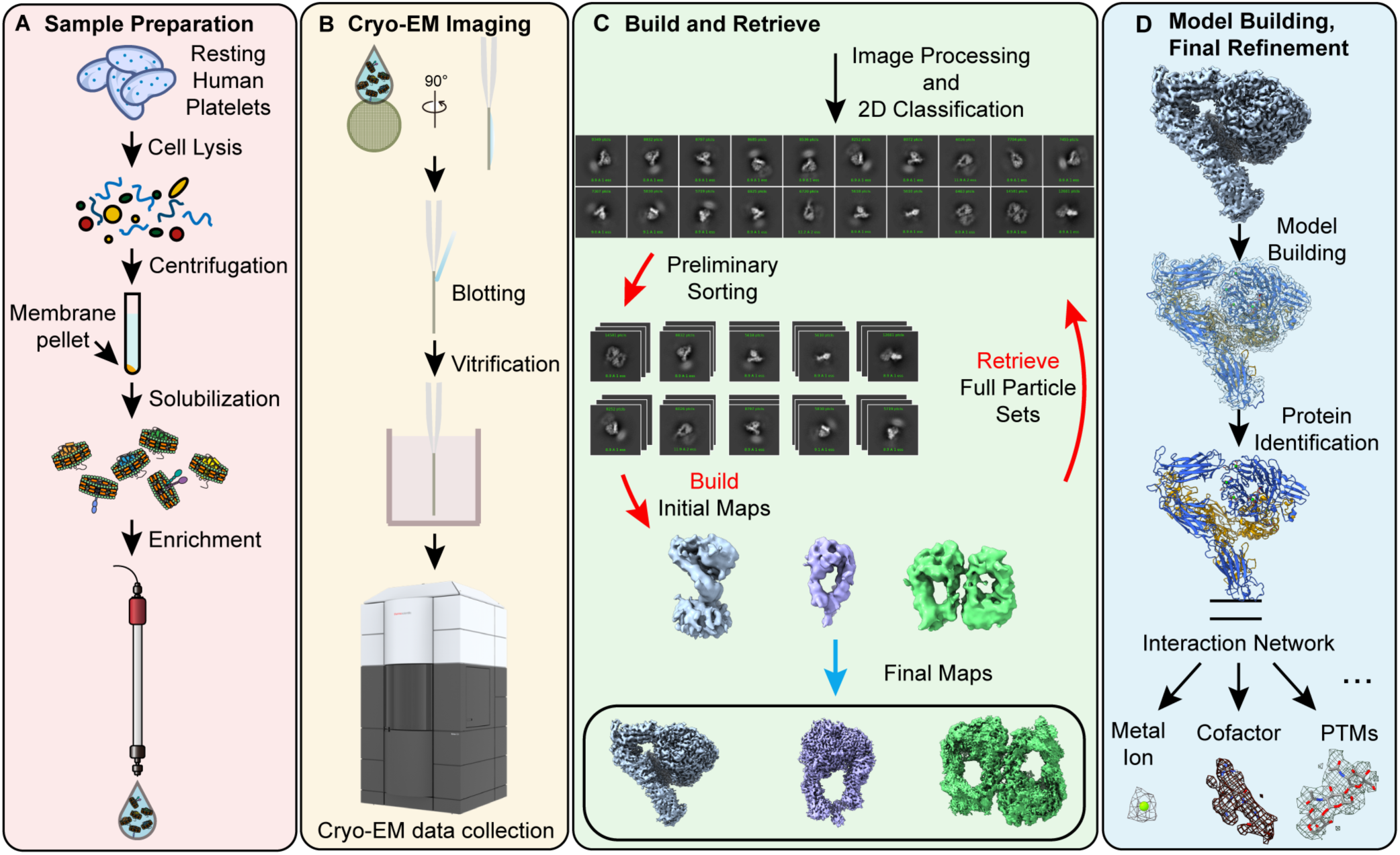
Build-and-Retrieve (BaR) Pipeline. Integrating cryo-EM with the BaR data processing methodology allows us to obtain atomic-level-resolution protein structures directly from crude preparations straight from their native environment.

In this paper, we applied cryo-EM coupled with BaR to resting platelet membrane proteins at ∼200 kDa from healthy individuals. We solved the structures of αIIbβ3 directly from native human platelet membranes. We identified αIIbβ3 structures in two distinct states, the inactivated and intermediate states, at 2.75 Å and 2.67 Å, respectively. We also observed a novel homodimer conformation at 2.85 Å in which the head regions of two αIIbβ3 molecules in intermediate states were facing each other.

## Materials and methods

### Cryo-EM sample preparation

Human platelet lysing, membrane protein extraction, and sample preparation for cryo-EM were performed as described previously. ^20,21^The detailed protocols for the preparation of human platelets and cryo-EM samples are described in the **Supplemental Methods**.

### Data collection and processing

A Titan Krios equipped with a K3 direct electron detector was used to collect data in super-resolution mode at 81,000 magnifications. Data were processed in the cryoSPARC V4.6.0 suite using the BaR protocol as described previously.^19–23^ The detailed data processing strategies are described in the **Supplemental Figure 1**. Protein Identification, model building, and refinement were performed as described.^19–23^ The detailed protocols for data collection, processing, model building and refinement are described in the **Supplemental Methods**.

### Quantification and statistical analysis

Standard Gold Standard-Fourier Shell Correlation curves at a threshold of 0.143 were computed using cryoSPARC v4.6.0 to obtain the final resolutions of protein models. The final atomic models were evaluated using MolProbity.

### Proteomic analysis of platelet membrane fractions

Sample preparation for proteomic analysis was performed as described previously. ^20,21^ The detailed protocols for protein digestion and identification are described in the **Supplemental Methods**.

## Results

Using cryo-EM coupled with the “Build-and-Retrieve” (BaR) data processing methodology (**Figure 1**), we solved three distinct cryo-EM maps at resolutions of 2.75 Å, 2.67 Å, and 2.85 Å, respectively (**Table 1** and **Supplemental Figure 1**). After model building and protein identification, all three maps were integrin αIIbβ3 in three different states: inactivated bent state, intermediate extended state, and a novel homodimer state that was formed by two integrin molecules in the extended state. Although we did not target integrin αIIbβ3 specifically or intentionally, it was not surprising that integrin αIIbβ3 was the first identified protein as it is one of the most abundant membrane proteins on human platelets at ∼200 kDa.^24^

**Table 1:**
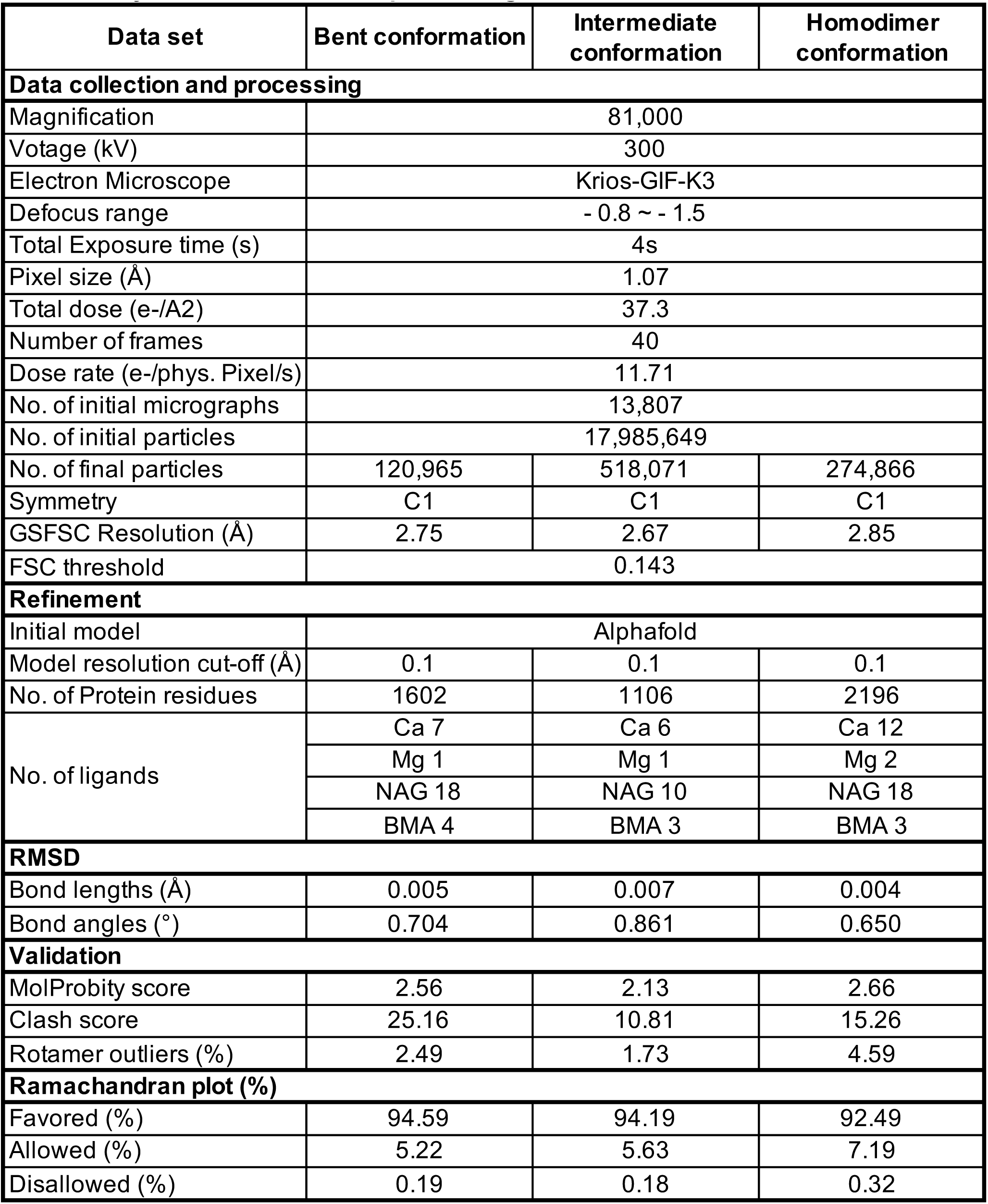
Cryo-EM data collection, processing, and refinement statistics.

We performed proteomic analysis to investigate the composition of the resting human platelet membrane peak at ∼200 kDa and to verify the existence of integrin αIIbβ3 in the sample. Our analysis identified approximately 800 different proteins within the peak. **Table 2** shows the 35 most abundant proteins in the peak, ranked by iBAQ (intensity-Based Absolute Quantification) values, which are calculated by summing the intensities of all detected peptides of a protein and dividing this sum by the number of theoretically observable peptides for that protein. We confirmed the presence of integrin αIIbβ3 in the sample (**Figure 2**, **Table 3**). Unsurprisingly, the integrin β3 subunit was the second most abundant protein, followed by integrin αIIb as the third (**Table 2**).

**Figure 2.**
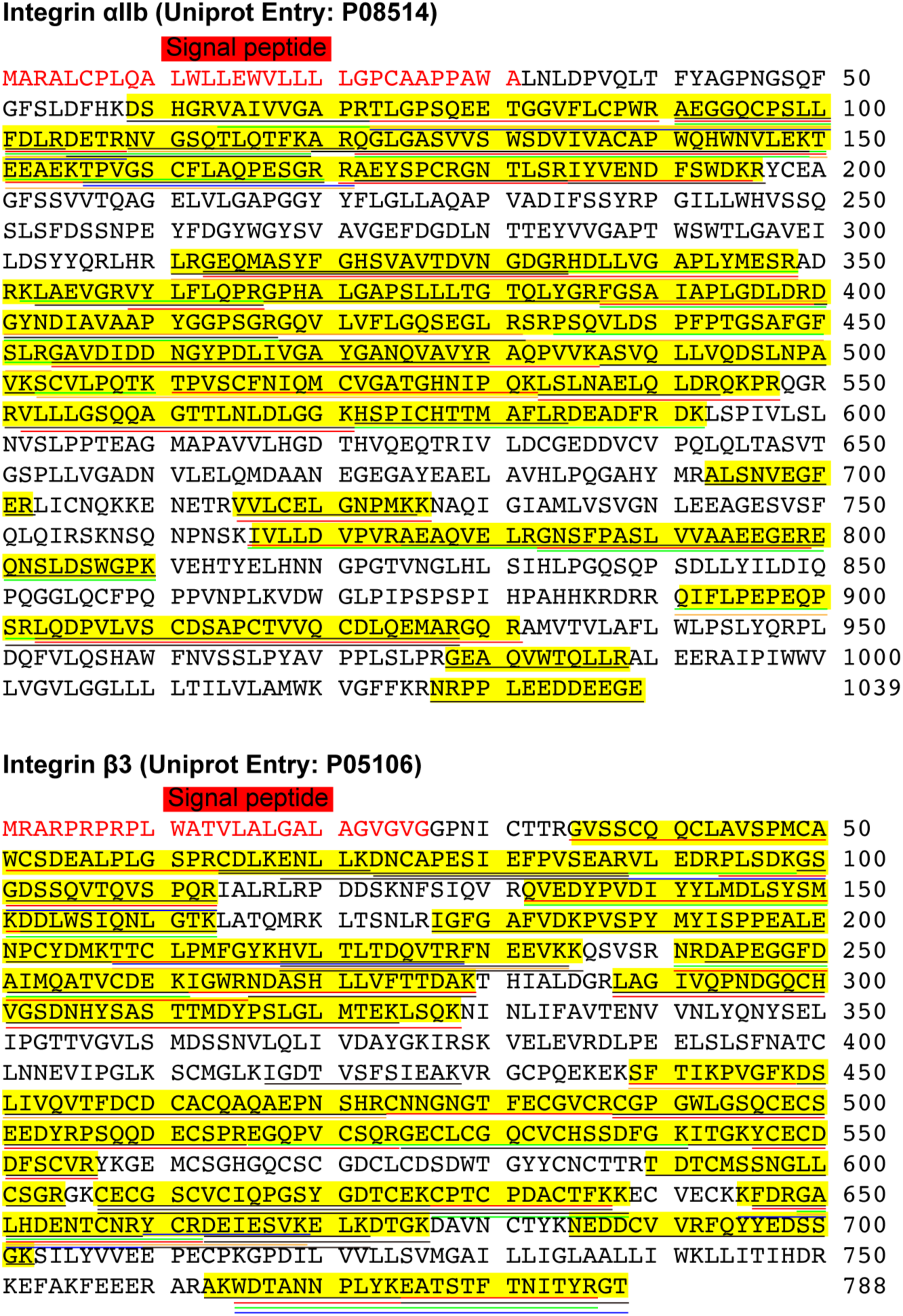
Peptide coverage of the human αIIbβ3 by MS. Full sequence of integrin αIIb (top) and β3 (bottom) were obtained from the UniProt database. The signal peptides were colored in red, and the mature proteins were colored in black. Each underline indicates a peptide of integrin αIIb (top) or β3 (bottom) identified from the mass spec study. Peptide coverage of integrin αIIb (top) and β3 (bottom) was highlighted in yellow.

**Table 2:** Top 35 proteins in the Mass Spec Study.

| **Protein IDs (Uniprot)** | **Protein names** | **Gene names** | **Mol. weight [kDa]** | **MS/MS count** | **iBAQ (High to Low)** | **Peptides** | **Sequence coverage [%]** |
| --- | --- | --- | --- | --- | --- | --- | --- |
| P02647 | Apolipoprotein A-I | APOA1 | 30.777 | 324 | 28.08583 | 42 | 83.1 |
| P05106 | Integrin beta-3 | ITGB3 | 87.057 | 1076 | 28.03847 | 59 | 63.8 |
| P08514 | Integrin alpha-lib | ITGA2B | 113.38 | 1856 | 26.63339 | 64 | 52.6 |
| P27105 | Erythrocyte band 7 integral membrane protein | STOM | 31.73 | 172 | 26.43232 | 18 | 72.9 |
| P02776 | Platelet factor 4 | PF4 | 10.845 | 32 | 26.42886 | 6 | 36.6 |
| Q93084 | Sarcoplasmic/endoplasmic reticulum calcium ATPase 3 | ATP2A3 | 109.25 | 349 | 26.30055 | 56 | 48.9 |
| P16671 | Platelet glycoprotein 4 | CD36 | 53.053 | 111 | 26.13606 | 20 | 37.1 |
| P13224 | Platelet glycoprotein Ib beta chain | GP1BB | 21.717 | 21 | 26.11728 | 6 | 23.8 |
| P21926 | CD9 antigen | CD9 | 25.416 | 28 | 26.07696 | 4 | 15.4 |
| P14406 | Cytochrome c oxidase subunit 7A2, mitochondrial | COX7A2 | 9.3959 | 11 | 25.94474 | 3 | 27.7 |
| P09669 | Cytochrome c oxidase subunit 6C | COX6C | 8.7813 | 10 | 25.89647 | 5 | 38.7 |
| P02675;C<br>ON__P02<br>676 | Fibrinogen beta chain | FGB | 55.928 | 347 | 25.84125 | 48 | 76.8 |
| P06576 | ATP synthase subunit beta, mitochondrial | ATP5B | 56.559 | 314 | 25.5996 | 29 | 73.7 |
| P25705 | ATP synthase subunit alpha, mitochondrial | ATP5A1 | 59.75 | 238 | 25.54687 | 49 | 66.9 |
| P61224;A<br>6NIZ1 | Ras-related protein Rap-1b | RAP1B | 20.825 | 60 | 25.5407 | 13 | 67.4 |
| P14770 | Platelet glycoprotein IX | GP9 | 19.046 | 41 | 25.51761 | 6 | 30.5 |
| P02679 | Fibrinogen gamma chain | FGG | 51.511 | 215 | 25.49344 | 38 | 72 |
| P05556 | Integrin beta-1 | ITGB1 | 88.414 | 223 | 25.41561 | 43 | 47.6 |
| P62873;P<br>16520 | Guanine nucleotide-binding protein G(I)/G(S)/G(T) subunit beta-1 | GNB1 | 37.377 | 80 | 25.26481 | 19 | 73.2 |
| P56385 | ATP synthase subunit e, mitochondrial | ATP5I | 7.9331 | 27 | 25.21097 | 7 | 71 |
| P48047 | ATP synthase subunit O, mitochondrial | ATP5O | 23.277 | 63 | 25.20202 | 15 | 75.1 |
| P20674 | Cytochrome c oxidase subunit 5A, mitochondrial | COX5A | 16.762 | 26 | 25.16779 | 7 | 38 |
| P07359 | Platelet glycoprotein Ib alpha chain | GP1BA | 71.539 | 76 | 25.16177 | 15 | 21.3 |
| O75947 | ATP synthase subunit d, mitochondrial | ATP5H | 18.491 | 67 | 25.00774 | 14 | 84.5 |
| P30273 | High affinity immunoglobulin epsilon receptor subunit gamma | FCER1G | 9.6674 | 8 | 24.9812 | 3 | 30.2 |
| P04899;P11488;P19087;A8MTJ3;P09471 | Guanine nucleotide-binding protein G(i) subunit alpha-2 | GNAI2 | 40.45 | 148 | 24.9715 | 29 | 72.4 |
| P16284 | Platelet endothelial cell adhesion molecule | PECAM1 | 82.521 | 248 | 24.93656 | 40 | 53.5 |
| Q9Y490;Q9Y4G6 | Talin-1 | TLN1 | 269.76 | 820 | 24.70055 | 156 | 73.4 |
| P02489 | Alpha-crystallin A chain | CRYAA | 19.909 | 54 | 24.5251 | 7 | 36.4 |
| P14625;Q58FF3 | Endoplasmin | HSP90B1 | 92.468 | 172 | 24.4853 | 43 | 53.1 |
| O75964 | ATP synthase subunit g, mitochondrial | ATP5L | 11.428 | 23 | 24.46251 | 6 | 52.4 |
| P23229 | Integrin alpha-6 | ITGA6 | 126.6 | 233 | 24.35824 | 56 | 50 |
| Q9Y277 | Voltage-dependent anion-selective channel protein 3 | VDAC3 | 30.658 | 53 | 24.32423 | 14 | 62.2 |
| P40197 | Platelet glycoprotein V | GP5 | 60.958 | 134 | 24.23783 | 24 | 50.5 |
| P21333;Q14315;O75369 | Filamin-A | FLNA | 280.74 | 658 | 24.2292 | 155 | 61.5 |

**Table 3:**
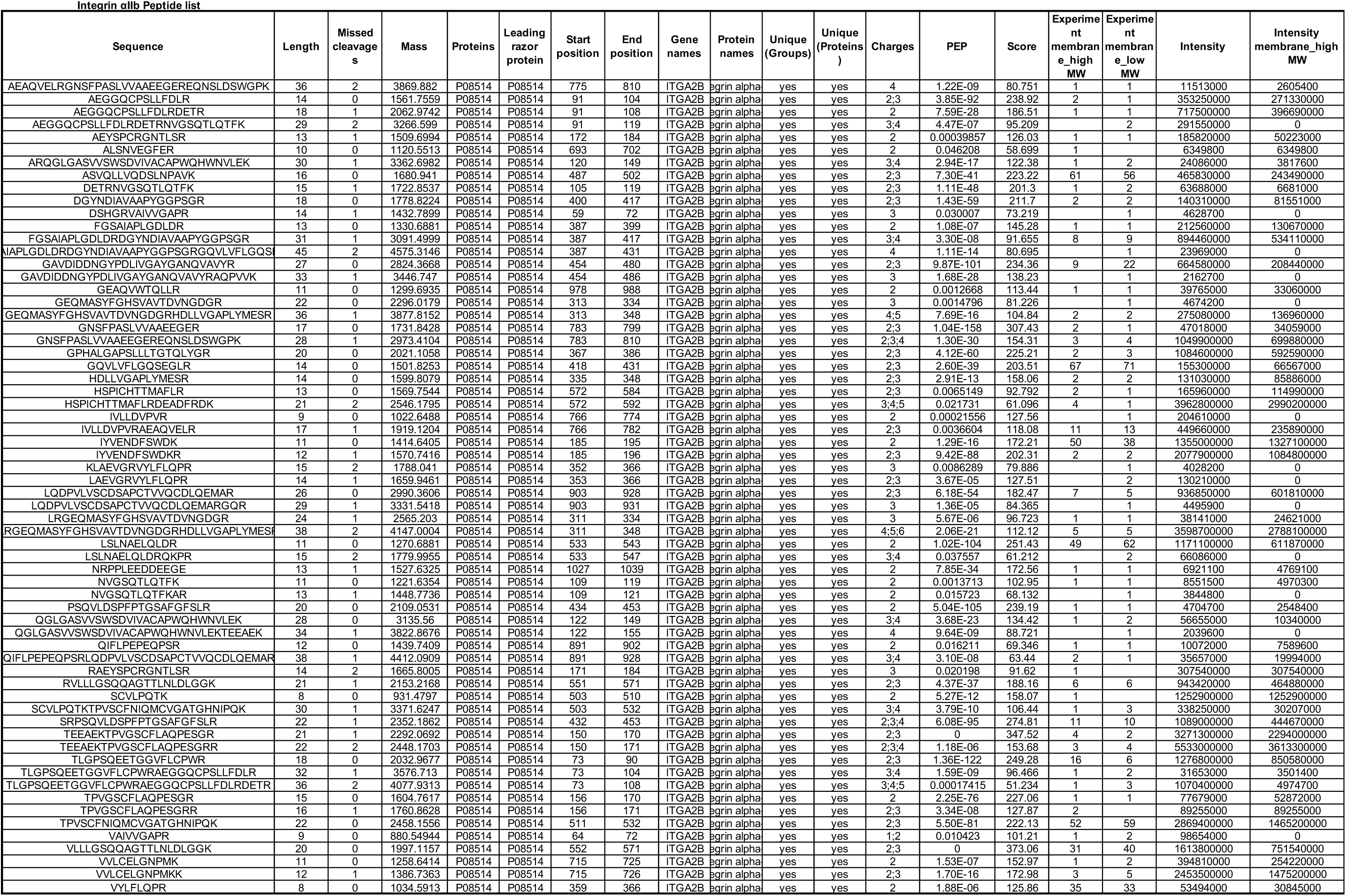
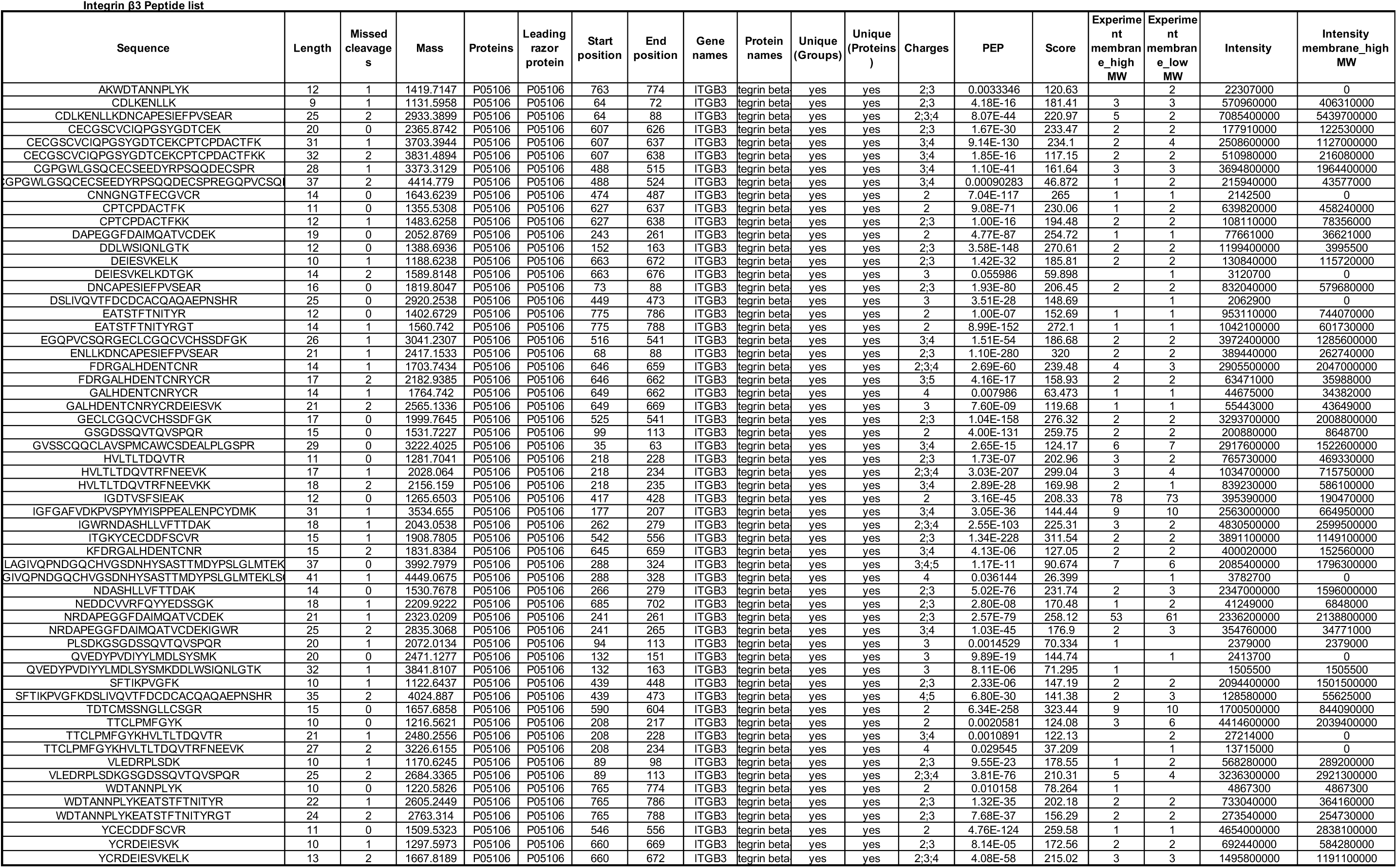
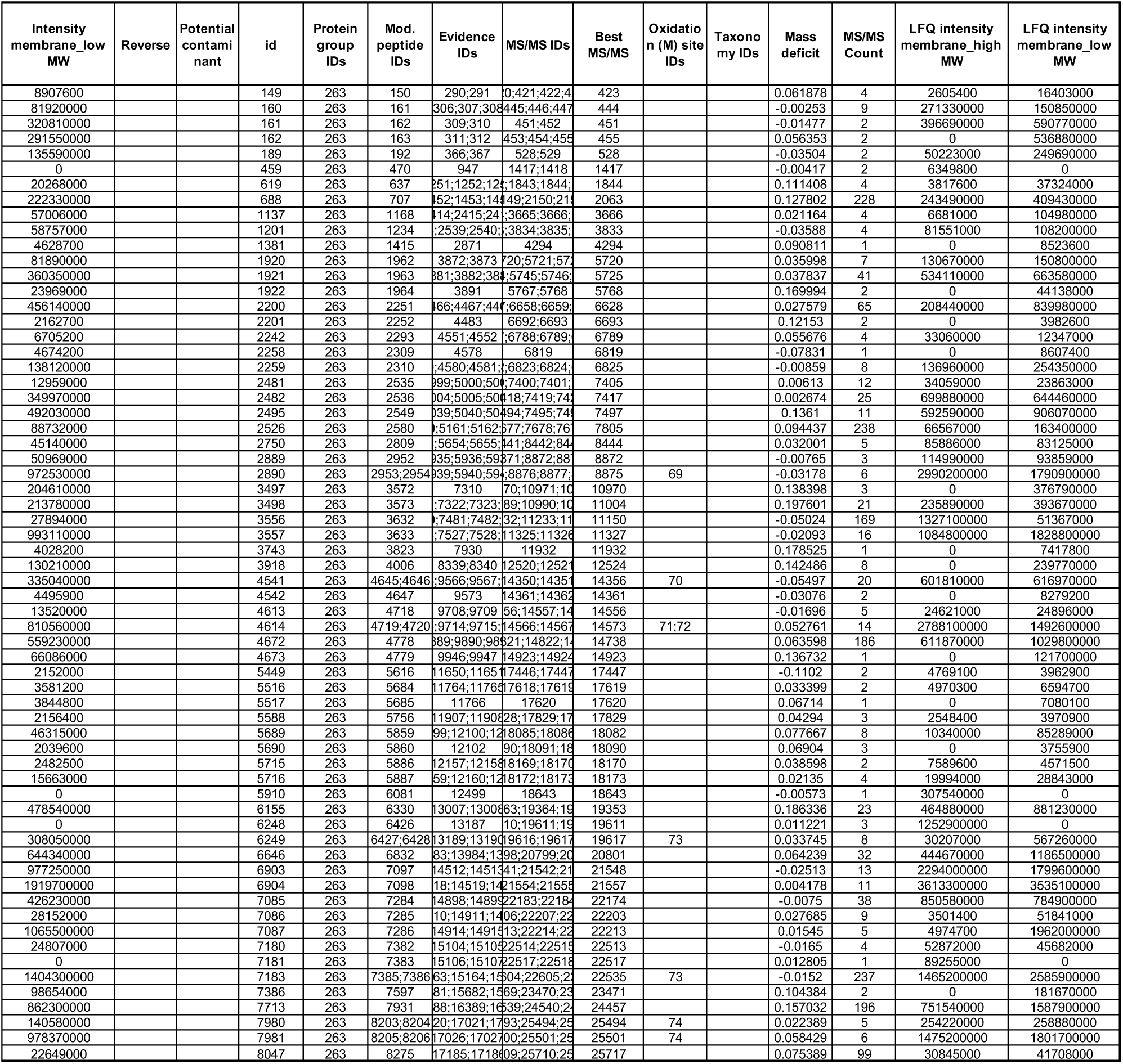
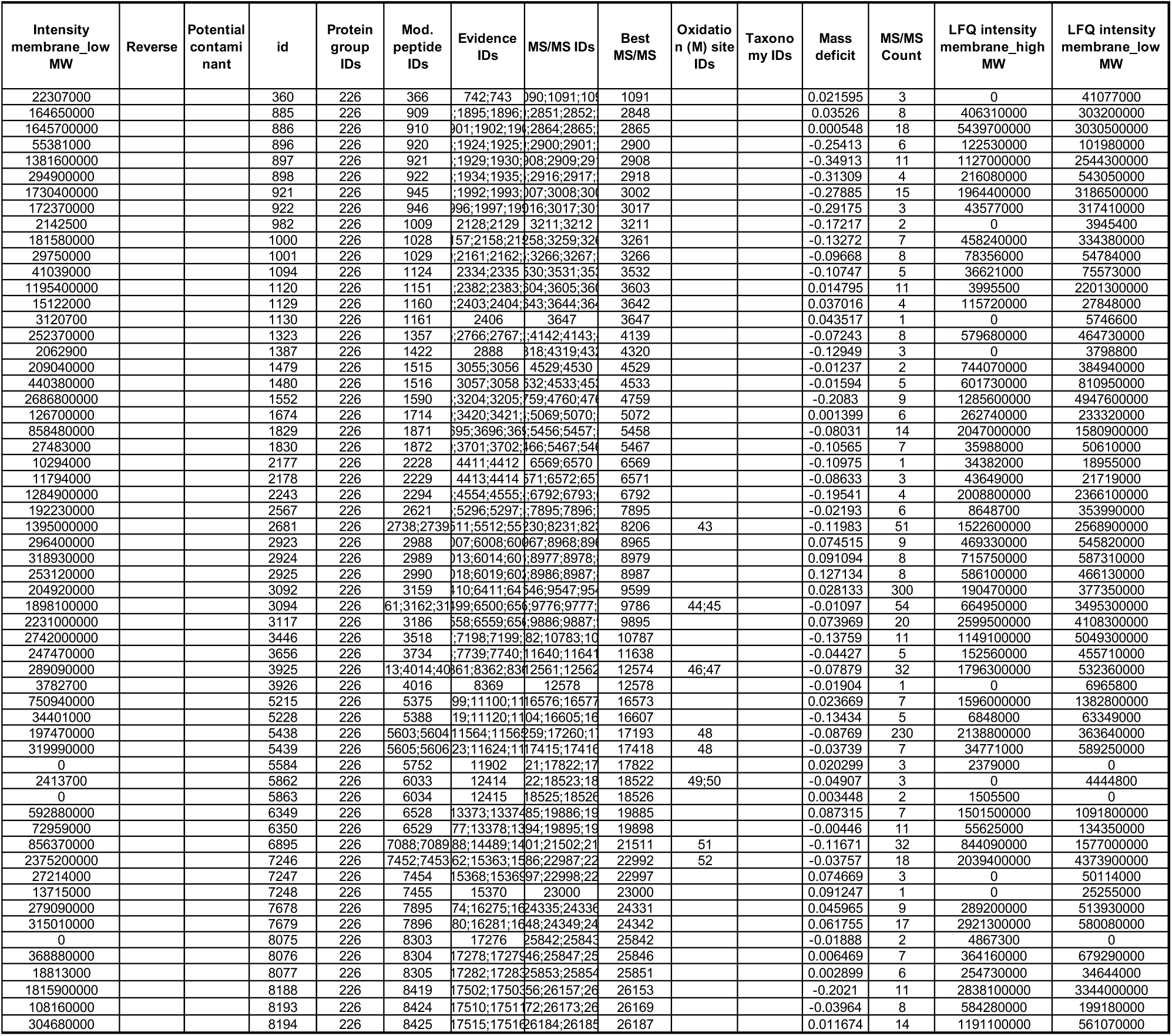
Peptide identified for ITGA2B and ITGB3 Integrin αIIb Peptide list.

### Inactivated bent state of αIIbβ3 - The overall architecture

The cryo-EM structure of native inactivated αIIbβ3 adopts the bent conformation seen previously in the crystal structures of the ectodomain, and the two most recent cryo-EM models of full-length integrin in native lipids and detergent, respectively.^8,10,25^ (**Figure 3, Supplemental Figure 2**) Our inactivated αIIbβ3 cryo-EM structure indicates that it is a heterodimer with overall dimensions of 114 Å x 82 Å x 110 Å, excluding the density of the nanodisc. The full topology of the heterodimer with all 12 extracellular subdomains of the ectodomain and the nanodisc density is clearly visible in the cryo-EM map, which makes assigning the molecule’s orientation to the membrane surface feasible. However, no density is detected for the transmembrane α-helices and αIIb and β3 short cytoplasmic tails. Our αIIbβ3 structure is superimposable onto the X-ray structure of complete ectodomain of integrin αIIbβ3 (3FCS.pdb^25^) with an RMSD between the respective 1,521 Cαs of 2.033 Å, and to the two recent inactive αIIbβ3 cryo-EM structures (8T2V.pdb^8^, and 8GCD.pdb^10^) with RMSDs between the respective 1,393 Cαs of 1.570 Å (8T2V.pdb) and 1308 Cαs of 1.141 Å (8GCD.pdb).^8,10^ The overall architecture of our integrin structure agree with the previous structural observation,^8,10,25,26^ indicating that solving high-resolution membrane protein structure directly from platelet membranes without purification using BaR data processing is feasible and valid.

**Figure 3.**
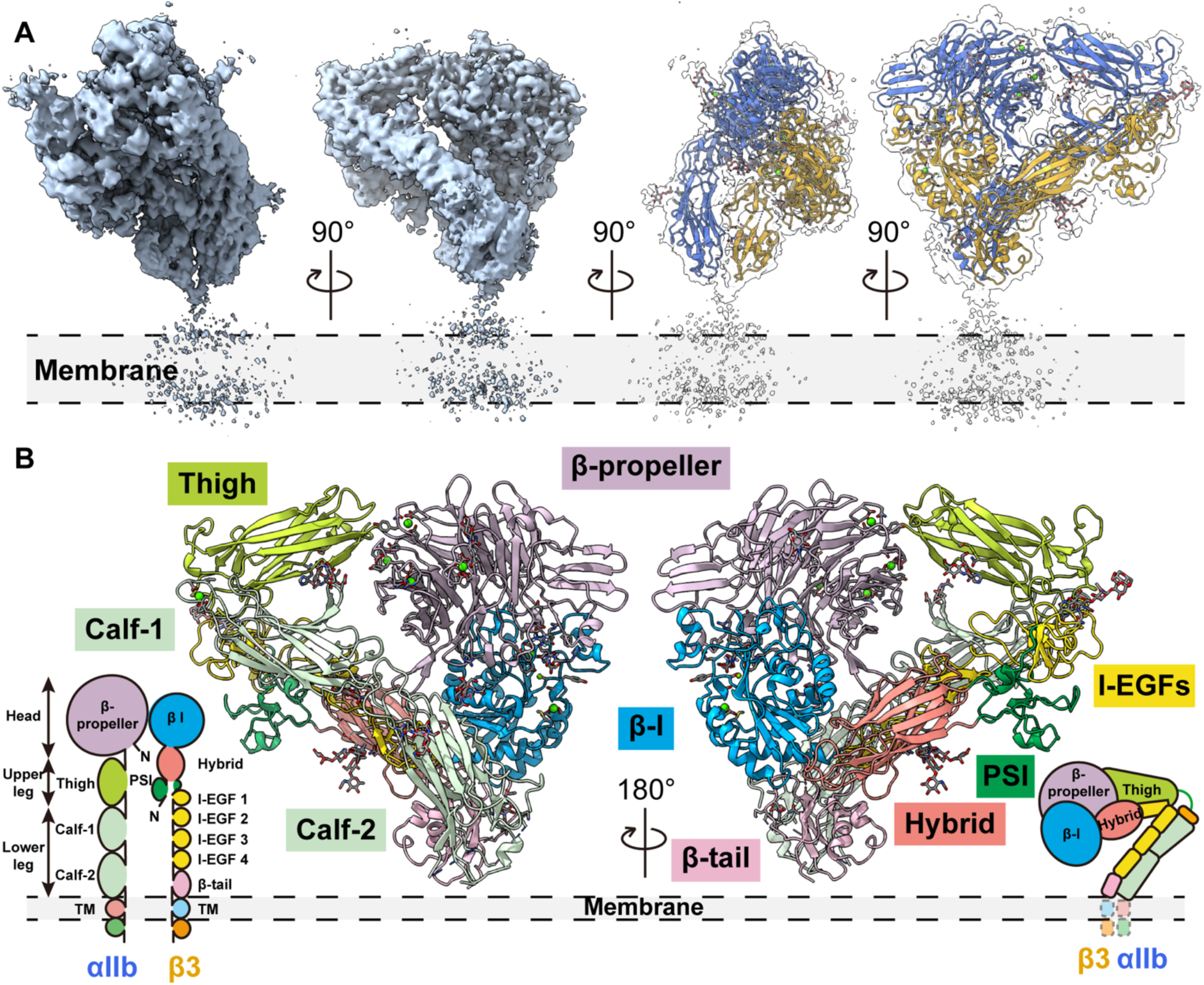
The overall architecture of integrin αIIbβ3 in the bent conformation. **A.** The overall cryo-EM map of integrin αIIbβ3 in bent conformation at 2.75 Å resolution is shown. The integrin αIIbβ3 model was shown in ribbon diagrams with the αIIb subunit colored in royal blue and the β3 subunit colored in goldenrod here and in subsequent figures. **B.** A total 12 subdomains of the ectodomain are resolved. The integrin αIIbβ3 model was shown in ribbon diagrams with each subdomain colored as follows: β-propeller in purple, Thigh in lime green, Calf in light green, β-I in blue, Hybrid in salmon, PSI in green, I-EGFs in yellow, and β-tail in pink.

The head and upper leg regions are 114 Å long measured from Thr150 (UniProt Entry: P08514 numbering) from the β-propeller domain to Thr501 at the end of the thigh domain; the lower leg region is 98.5 Å long measured from the calcium at the αIIb genu point to Glu991 at the end of the calf-2 domain. Although the TM α-helixes are not solved in the bent αIIbβ3 conformation, we can still assign the orientation of the ectodomain to the plane of the membrane since the nanodisc density is clearly visible. (**Figure 3**) The molecule forms an overall “squatting” conformation. (**Figure 3**) The β-propeller domain of the αIIb subunit forms a 36.4° angle to the thigh domain, and a 48.1° angle formed between the thigh and calf-1 domains. The calf-2 domain in the lower leg region forms a 62.5° angle to the cell membrane. (**Figure 3, Supplemental Figure 2**) The head and upper leg regions of the β3 subunit (βΙ, hybrid, and PSI domains) form at a sharper 16.2° angle to the lower leg region (EGFs and β-tail domains), and the lower leg region forms a 36.1° angle to the cell membrane. (**Figure 3, Supplemental Figure 2**) The head and upper leg regions of αIIbβ3 form a plane that is tilted to the cell surface with a 49.2 ° angle, which leads to the more “relaxed” bending on the αIIb subunit and a tighter bending conformation on the β3 subunit. (**Figure 3, Supplemental Figure 2**) The molecule is 110 Å in height, measured from the highest point of the head region (His61 from the β-propeller domain) to the membrane surface. (**Figure 3, Supplemental Figure 2**)

Using the 3D Flexible Refinement (3DFlex) function embedded in the CryoSparc v4.6.0 suite^27,28^, we determined the local motion and dynamics of the flexible αIIbβ3 from the cryo-EM dataset in the bent conformation. (**Supplemental Figure 2E**, and **Movie S1**) Even in the single high-resolution “canonical” 3D density map of αIIbβ3, the molecule displays conformational variability and flexible (nonrigid) motion: the lower leg region demonstrated the highest dynamics followed by the upper leg region. (**Supplemental Figure 2E**, and **Movie S1**) This flexibility leads to the molecule continuously oscillating on the cell membrane. In contrast, the entire head region is quite stable with minimal dynamics being observed. This is also in agreement with our local resolution map (**Supplemental Figure 2F**): the head region has the highest resolution (∼2.5 Å) followed by the upper leg region (∼3 Å). Due to the molecular flexibility, the resolution of the lower leg region ranges from 3.5 Å to +5 Å, where the β-tail has the lowest resolution.

### Inactivated bent state of αIIbβ3 - Glycosylation and ion binding sites

As a glycoprotein, integrin αIIbβ3 has a total of eleven predicted N-linked glycans: five on the αIIb subunit (αIIb-N46 and αIIb-N280 in the β-propeller domain, αIIb-N601 in the thigh domain, αIIb-N711 in the calf-1 domain and αIIb-N962 in the calf-2 domain) (UniProt Entry: P08514 numbering); and six on the β3 subunit (β3-N346 in the βI domain, β3-N125 and β3-N397 in the hybrid domain, β3-N478 in the I-EGF1 domain, β3-N585 in the I-EGF3 domain and β3-N680 in the β-tail domain) (UniProt Entry: P05106 numbering). ^29^ Clear densities of all 11 N-glycans are visible in our cryo-EM map of integrin αIIbβ3, which was directly extracted from resting human platelet membranes without manipulation during sample enrichment (**Figure 4A**). In agreement with previous structural data,^30^ we observed all ion binding sites on the molecule (**Figure 4B**). In other structural work that focused on purified αIIbβ3, additional Mg^2+^, Ca^2+^, or both were added to the buffer throughout the purification process. In contrast, we did not provide exogenous divalent ions to our heterogenous cryo-EM sample during our sample preparation except at the initial washing step of the intact platelet membrane with high salt buffer to remove soluble proteins. Therefore, all ion binding was straight from the original whole blood sample from the donors. There are four Ca^2+^ binding sites at the β-propeller domain, one Ca^2+^ at the αIIb genu point, and three divalent-ions at the βI-domain: a Mg^2+^ at the metal ion-dependent adhesion site (MIDAS), a Ca^2+^ at the adjacent to MIDAS (ADMIDAS), and a Ca^2+^ at the synergistic metal ion-binding site (SyMBS). (**Figure 4B**)

**Figure 4.**
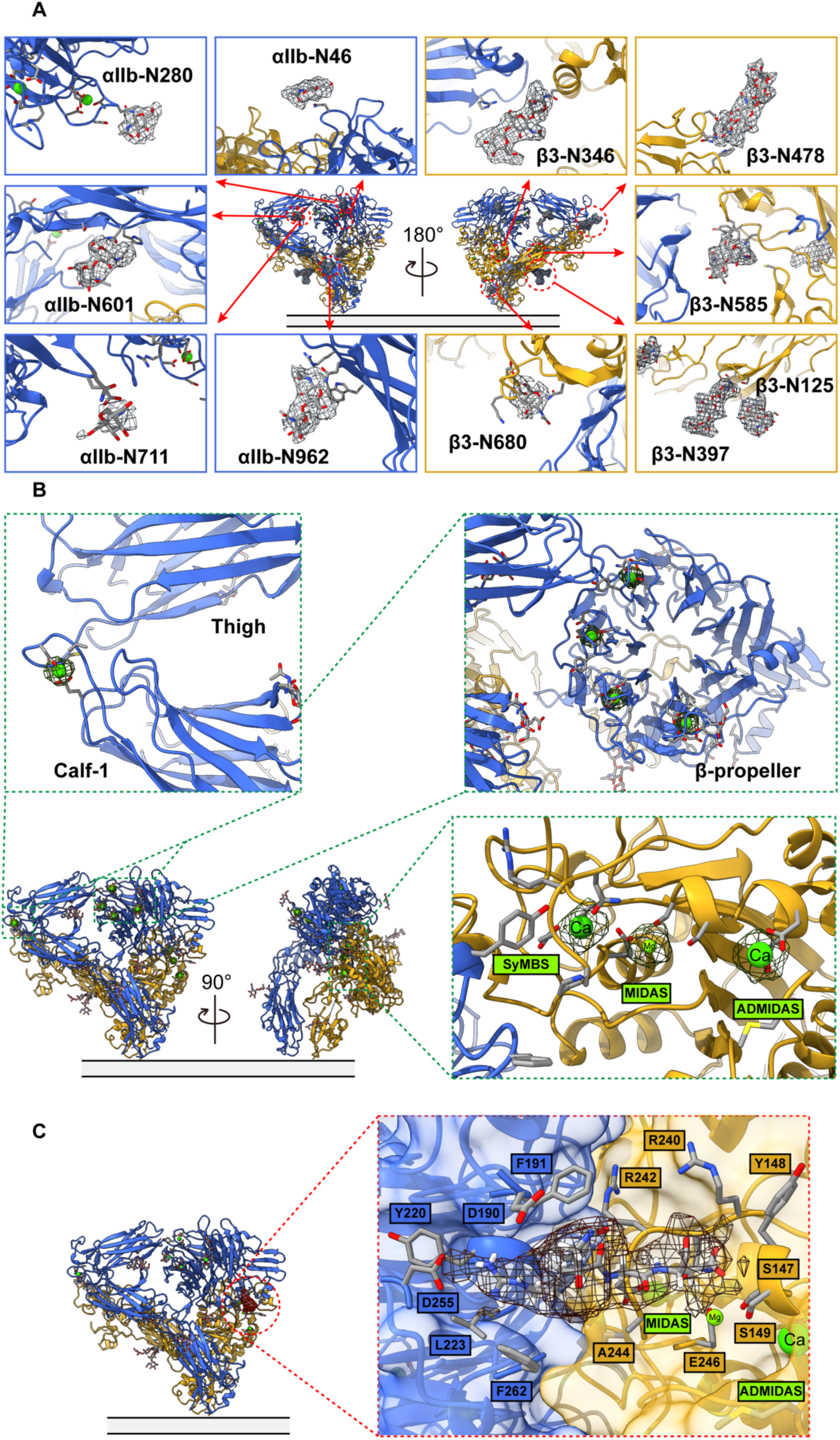
The interaction network of integrin αIIbβ3 in the bent conformation. **A.** 11-N-linked glycan sites were preserved in the bent integrin αIIbβ3 obtained from the native environment**. B.** Closeup of the cryo-EM density of the metal ions, one at the α-genu (top left insert), four at the β-propeller (top right insert), and three at SyMBS (occupied by a Ca^2+^), MIDAS (occupied by a Mg^2+^), and ADMIDAS (occupied by a Ca^2+^) of the β-I domain (bottom right insert). **C.** A clear density of an RGD-motif-containing peptide was observed in the ligand binding site.

### Inactivated bent state of αIIbβ3 - Liganded by an RGD-motif-containing peptide

Interestingly, the bent inactivated αIIbβ3 structure is liganded by a long peptide that contains an Arg-Gly-Asp (RGD)-motif. (**Figure 4C**). In this “squatting” conformation, the fully accessible ligand binding site at the heterodimer interface of αIIbβ3 is facing forward and is 62 Å from the membrane surface. The accessibility of the ligand binding site of αIIbβ3 in its bent conformation agrees with the two most recent αIIbβ3 cryo-EM structures.^8,10^

In our cryo-EM density map at 2.75 Å resolution, the ligand binding site of bent inactivated αIIbβ3 is occupied by a clear density of an RGD-motif-containing peptide. Our αIIbβ3 structure is superimposable onto the X-ray structure of integrin αIIbβ3 headpiece bound to fibrinogen γ chain peptide by Springer and colleagues^26^ (2VDO.pdb) with an RMSD between the respective β-I domain 168 Cαs of 0.366 Å, an RMSD between the β-propeller domain 390 Cαs of 0.258 Å, and an RMSD between the entire headpiece 658 Cαs of 0.729 Å. In agreement with the previous work, the surface representation of the binding site shows that the RGD peptide sits on the grove formed between the heterodimer interface of αIIbβ3. Since our BaR sample of resting human platelet membrane was a direct extraction from the endogenous resource, and the RGD-motif is a conserved sequence recognized by the integrin binding site, this RGD-containing peptide should belong to a natural ligand of αIIbβ3. We were able to observe some clear ligand density extended from the RGD-motif (**Supplemental Figure 2H**). A long density extended out from the αIIbβ3 binding pocket that almost act as a staple to tape the molecule in its bent conformation (**Supplemental Figure 2H**). However, without knowing the precise amino acid sequence, we were unable to identify or track the origin of the peptide due to the weak electron density of the peptide.

### Intermediate state of αIIbβ3 - The overall architecture

The highly dynamic nature of αIIbβ3 is required to fulfill its role in mediating inside-out and outside-in signaling. We also observed a second conformation of αIIbβ3 is the intermediate state in our BaR dataset from resting human platelet membrane. These structures indicate that the protein also displayed a flexible structural rearrangement on the inactivated platelet. Although the lower leg region was visible in the 2D classification (**Supplemental Figure 3B**), the electron density of the lower leg region was lost during the following data processing due to the highly dynamic nature of the protein. Only the closed headpiece, which contained the full head region and the upper leg region (β-propeller, Thigh, and partial Calf-1 domains on the αIIb subunit; and βI, Hybrid, PSI, and partial I-EGF1 domains on the β3 subunit) in the intermediate state of αIIbβ3 at 2.67 Å was solved. (**Figure 5A, Supplemental Figure 3**).

**Figure 5.**
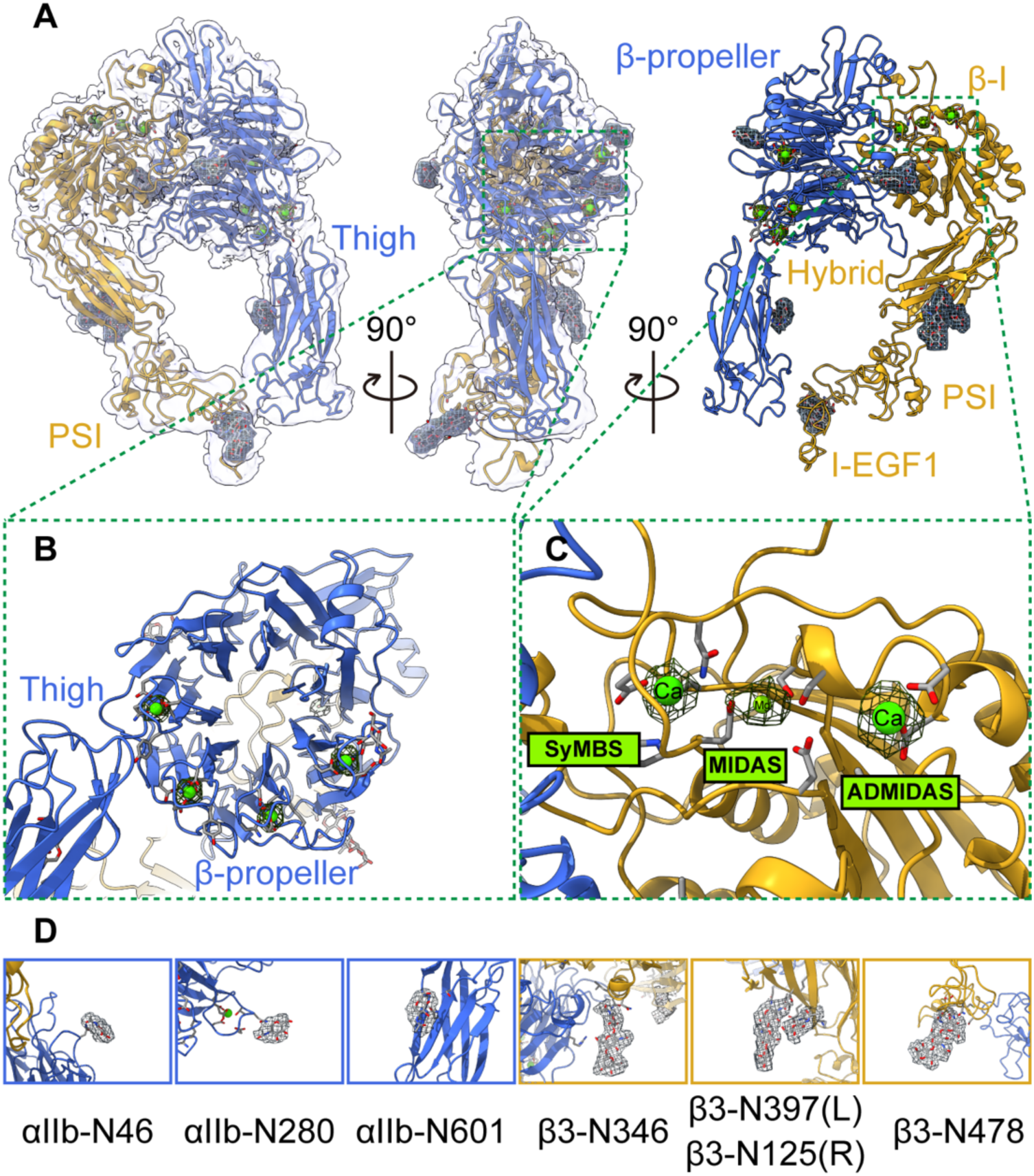
The architecture of integrin αIIbβ3 in the intermediate conformation and its interaction network. **A.** The overall cryo-EM map of integrin αIIbβ3 in intermediate conformation at 2.67 Å resolution is shown. The right panel is the side of the headpiece facing the cell membrane. **B.** Closeup of the cryo-EM density of the four metal ions at the β-propeller domain. **C.** Closeup of the cryo-EM density of the three metal ions at SyMBS (Ca^2+^), MIDAS (Mg^2+^), and ADMIDAS (Ca^2+^) of the β-I domain. D. 7-N-linked glycan sites within the solved region of the intermediate integrin αIIbβ3 were preserved.

### Comparison between intermediate and bent states of αIIbβ3

There is a distinct difference between the top views of the bent and intermediate states, which validated there were indeed two different conformations of integrin αIIbβ3 in our dataset. The resolution of the density maps of both bent and intermediate states at the ligand binding site was around 2.5 Å. At this resolution, we can confidently assign all side chains of each amino acid. In contrast to a clear electron density of an RGD-motif-containing peptide in the bent conformation, the ligand-binding site of the intermediate state was unoccupied, which convinced us to separate those particles into two different states. In addition, the density of three divalent ions at the MIDAS, ADMIDAS, and SyMBS were equally strong in the bent conformation. (**Figure 4B**) However, the electron density of the Mg^2+^ at the MIDAS was significantly weaker than the other two Ca^2+^ in the intermediate state. (**Figure 5C**) The Mg^2+^ at the MIDAS is critical for αIIbβ3 ligand binding, which further supports that the closed headpiece in the intermediate state is unliganded. Further, an additional density can be observed in the 2D projection of the top view of the bent conformation but not in the intermediate state. This density came from the lower leg region, which can be seen through the center of the headpiece from the top of the molecule. (**Supplemental Figure 4A**) On the other hand, the lower leg region extends outward and is no longer visible through the center of the headpiece in the intermediate state. (**Supplemental Figure 4A**) The RMSD between the respective 834 Cαs of the intermediate state and the same headpiece region in our inactivated bent state is 1.662Å, which indicates these two conformations are similar but not the same. Compared to the inactivated state, the most noticeable overall structural shift of the headpiece came from the thigh domain in the αIIb subunit (shifted 9 Å) and the hybrid (shifted 6 Å), PSI, and I-EGF1 domains (shifted 9 Å) in the β3 subunit. (**Supplemental Figure 4B**).

Seven glycosylation sites were elucidated within the solved region of the intermediate state: three on the αIIb subunit (αIIb-N46 and αIIb-N280 in the β-propeller domain and αIIb-N601 in the thigh domain); and four on the β3 subunit (β3-N346 in the βI domain, β3-N125 and β3-N397 in the hybrid domain, and β3-N478 in the I-EGF1 domain). This is similar to the inactivated state. (**Figure 5D**)

### Novel homodimer of αIIbβ3 - The overall architecture and dimer interface

A novel homodimer conformation of αIIbβ3 at 2.85Å was also observed in which the head regions of two αIIbβ3 molecules in intermediate states were facing each other. (**Figure 6, top, Supplemental Figure 6**) This additional homodimer interface was formed between two αIIb subunits and locked two molecules in a “face-to-face” and “head-to-toe” conformation. (**Figure 6, bottom**) Specifically, there are two clusters of interaction formed between Ser60 - Gly62 on the β-propeller domain and Pro574 - Cys576 on the thigh domain, and another two clusters of interaction formed between Gly570 – His572 on the thigh domain and Asp460 on the β-propeller domain

**Figure 6.**
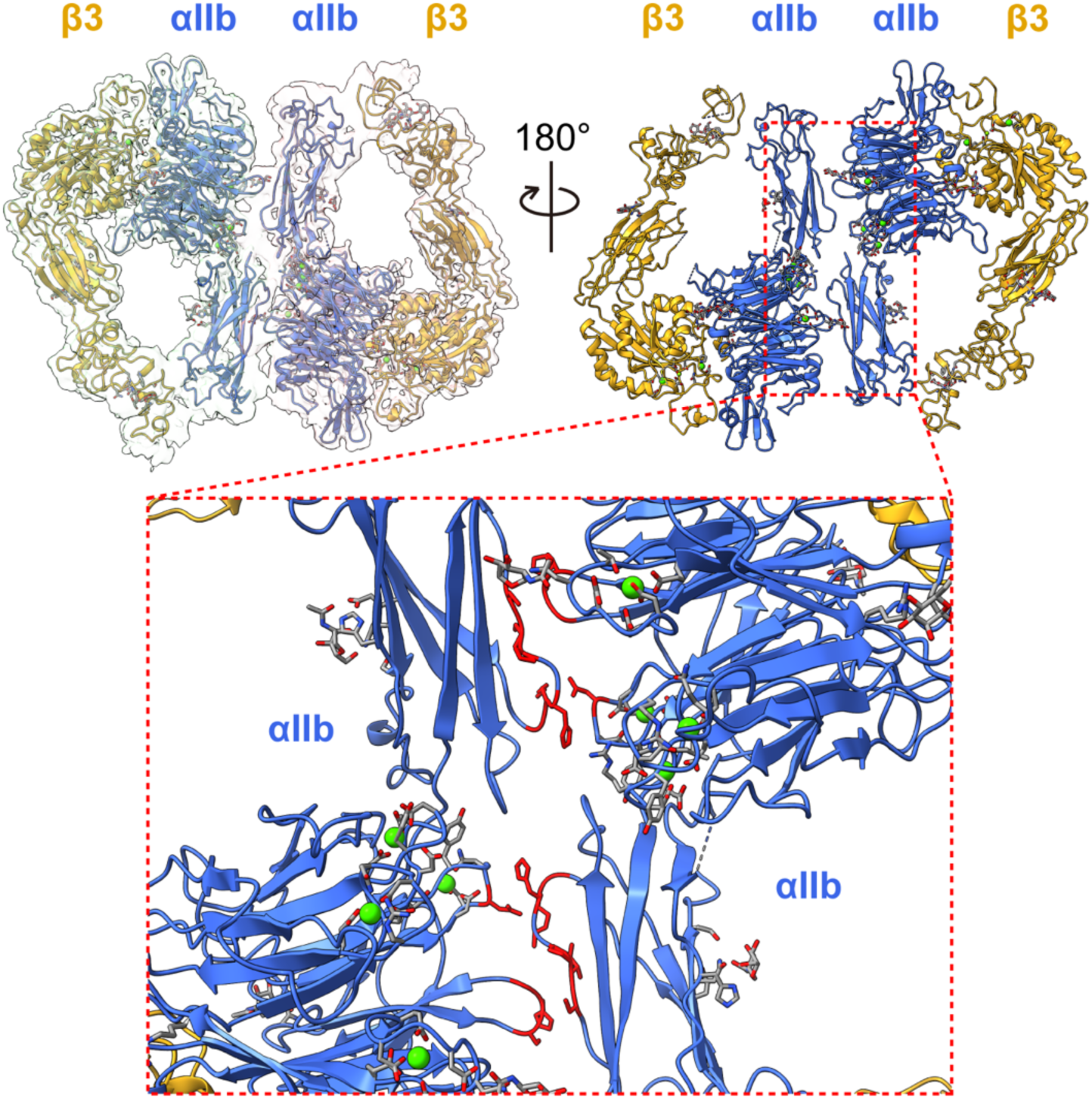
A novel homodimer conformation formed by two intermediate integrin αIIbβ3 molecules was revealed. The overall cryo-EM map of integrin αIIbβ3 in homodimer conformation at 2.85 Å resolution is shown. The top right panel is the side of the headpiece facing the cell membrane. The bottom insert shows the closeup of the dimer interface formed by two αIIb subunits from two individual integrin αIIbβ3 molecules. The residues that are involved in the dimer interface were colored in red.

The homodimer showed similar glycosylation and ion binding profiles compared to the previous two structures. Like the intermediate conformation, we did not observe any ligand density in either αIIbβ3 molecule of the homodimer. (**Figure 6, top**) Together with our observations in the first two structures, this unique integrin αIIbβ3 architecture identified directly from the native platelet membrane may indicate a previously unknown self-regulatory effect of αIIbβ3 to temporally regulate its ligand accessibility in the intermediate state.

## Discussion

Platelets are essential for the vascular system’s integrity and primary hemostasis. To keep the perfect balance between the quiescent inactivated state in normal conditions and quick activation upon injury, platelet function is orchestrated by a network of interactions between proteins and biomacromolecules on the membrane to sense and relay the extracellular stimulus across the membrane.^1,2,7^ Mutations, structural modifications, or changes in expression levels of these proteins can profoundly affect platelet physiology and are associated with numerous disorders. ^3,31^ Further, the native membrane environment provides essential additional regulatory cues, such as ions, post-translational modifications, and endogenous ligands *in situ* that coordinate the protein structure and mechanism of action. Defining the physical interactions of this network of membrane proteins in the endogenous environment will shed light on understanding basic platelet biology and physiology.

Technological breakthroughs in single-particle cryo-EM enable solving near-atomic-resolution structures of macromolecules without high-quality crystals. This is critical for studying membrane proteins and larger complexes, as their sample preparation and crystallization are difficult. The rapid developments of cryo-EM expand the molecular weight range and allow protein structures to be solved directly from native cell extracts. However, solving multiple proteins or multiple states of a single protein from a raw heterogeneous sample remains a challenge for single-particle cryo-EM.

In this study we combined cryo-EM with the Build-and-Retrieve (BaR) data processing method to analyze the platelet membrane proteins to bridge structural biology with -omics. BaR is an iterative bottom-up approach that uses *in silico* purification to obtain high-resolution cryo-EM maps to solve structures of proteins in different states in the native environment.^19–23^ Here, we report the unmodified integrin αIIbβ3 structure obtained directly from resting human platelet membranes in its inactivated and intermediate states at 2.75Å and 2.67Å, respectively. We also solved a novel homodimer conformation formed by two intermediate-states of αIIbβ3 at 2.85Å.

Integrin αIIbβ3, also known as GPIIb/IIIa, is a cell surface receptor that plays a crucial role in platelet function. When activated, αIIbβ3 undergoes conformational changes that enable it to bind to fibrinogen and other ligands, which further facilitates platelet aggregation. The previous structural studies with αIIbβ3 in different states have provided key insights on its mechanism of action. Our data with αIIbβ3 directly from resting human platelets add new information on the highly dynamic nature of this protein prior to activation. We also identified the structural rearrangement between the three states (**Figure 7**) and the local dynamics and flexibility of αIIbβ3 in the bent conformation.

**Figure 7.**
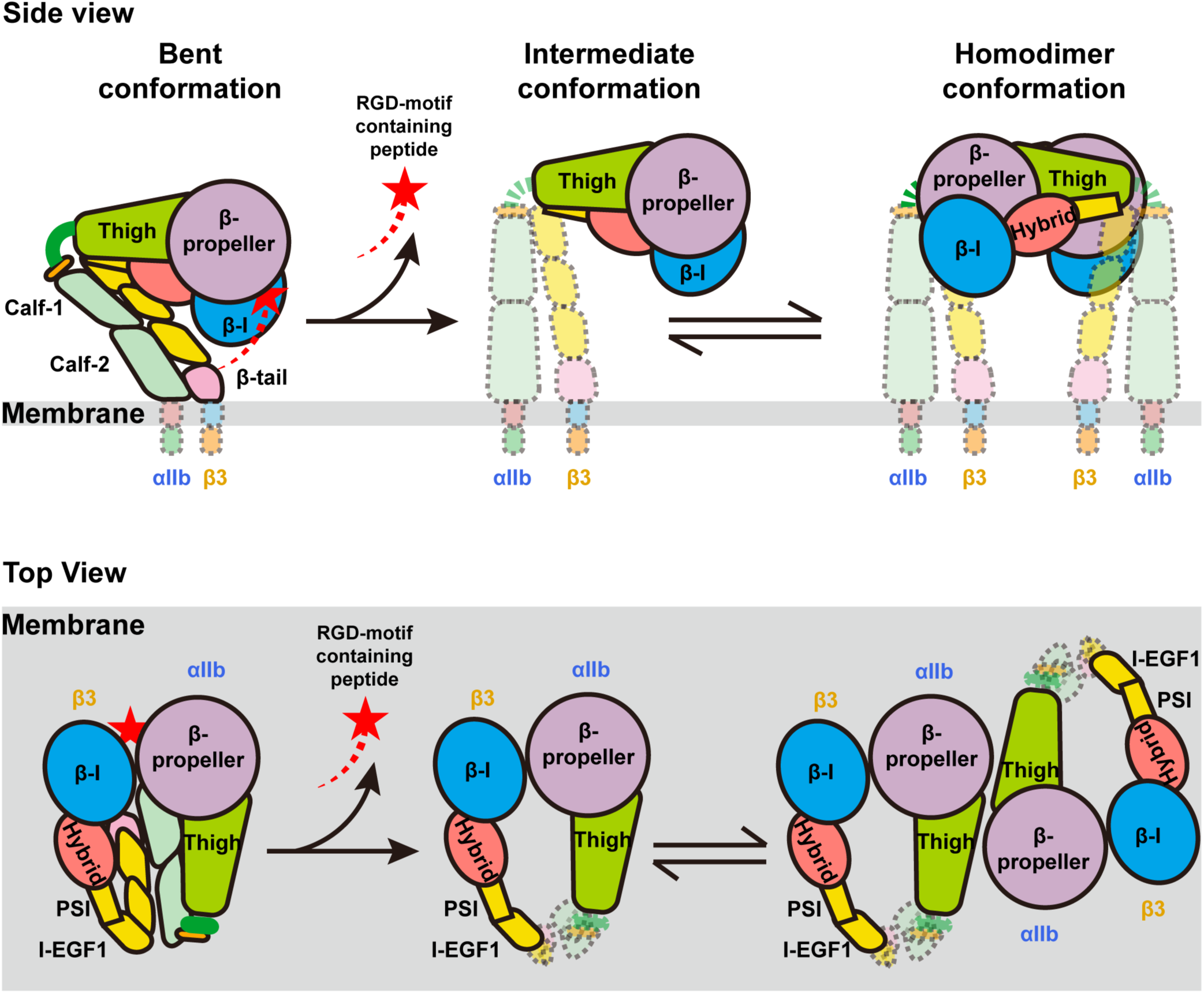
Proposed model for the dynamic of integrin αIIbβ3 on resting human platelet membrane. On the surface of the resting human platelets, integrin αIIbβ3 has three major conformations: inactivated bent, intermediate, and homodimer. An RGD-motif-containing ligand (red shooting star) binds to the bent conformation and keeps the ligand binding site occupied when the molecule is inactive. The ligand releases while αIIbβ3 extends to the intermediate state. Two adjacent αIIbβ3 molecules in the intermediate state can form a homodimer on the surface. There is a dynamical conformational equilibrium between the intermediate state and the homodimer state of αIIbβ3 on the human platelets. The top panel shows the side view of the molecules in which the extracellular side is on the top and intracellular side is on the bottom. The bottom panel shows the top view of the molecules where the platelet membrane (grey) lays on the bottom.

Our αIIbβ3 structures in bent and intermediate conformations are in agreement with their corresponding crystal and cryo-EM structures. ^8,10,25^ Our cryo-EM sample was prepared directly from its native environment without additional affinity enrichment. This preserved the natural regulatory cues, such as glycosylation sites, ion binding, and co-factors, which new insights on how the environment regulates αIIbβ3 structure and function.

First, N-linked glycosylation is critical to regulating the adhesion function of integrins.^29^ The role of each N-linked glycan on αIIbβ3 integrin structure and function was thoroughly studied by mutagenesis in HEK293FT cells.^29^ Our structural data provided additional clear visual support for the presence of all 11 N-linked glycans. (**Figure 4A**) Micro-, macro-, and meta-heterogeneity of glycosylation can impact protein structure and function.^32^ We observed some level of micro- and macro-heterogeneity of N-linked glycosylation among the bent, intermediate, and dimer conformations. The structural and functional impact of heterogeneous glycosylation of integrin αIIbβ3 is an area for future exploration. The electron density was weaker in the I-EGF1 domain in both the intermediate and the dimer conformations. Thus, although we did observe a clear N-linked glycan around the N478 site in the β3 subunit, assigning the precise location of the NAG was challenging given that two additional asparagine residues (N475 and N509) were in close proximity. Future studies will determine the functional significance of the interplay of these three asparagines on N-linked glycosylation in the I-EGF1 domain.

Second, we observed all ion binding sites on the αIIbβ3 molecule. (**Figure 4B**, **5B** and **5C**) Our goal is to solve structures of proteins from the human platelet membranes with minimal manipulation. We did not provide additional divalent ions during our sample preparation other than in the high salt buffer for removing soluble proteins from the intact platelet membrane at the initial step. To our surprise, there were clear electron densities of four Ca^2+^ binding sites at the β-propeller domain, one Ca^2+^ at the αIIb genu point, and three divalent-ions binding at the MIDAS, ADMIDAS, and SyMBS at the βΙ-domain, which indicated all ion binding was from the original whole blood from the donor. This demonstrates the power of our approach to preserve the interactions from the original native environment.

Third, the αIIbβ3 cryo-EM map of the bent conformation showed a clear density from a peptide that contains an RGD-motif at the ligand binding site. (**Figure 4C**) This agrees with the overall architecture of the bent conformation and the most recent cryo-EM structure of the full-length αIIbβ3,^8^ in which the whole molecule was “squatting” on the lipidic nanoparticles and the ligand binding site was fully accessible. This could potentially challenge the conventional αIIbβ3 switchblade model on its activation mechanism, in which the headpiece of the molecule points to the membrane to maintain its low affinity for the ligand.^8^ We cannot rule out that this observation may be due to the sample extraction strategy. Coller suggested that extraction conditions, by detergent only or by inserting into nanodiscs, affected the overall structure of αIIbβ3.^33^ Therefore, further studies are needed to carefully evaluate the impact of the lipidic mimetic environment on αIIbβ3 structure.

We also identified a third conformation, which is a novel homodimer of αIIbβ3. (**Figure 6**) This new αIIbβ3 conformation highlights the strength of the BaR method in analyzing crude samples straight from their original sources with minimum manipulation in protein purification. Unlike other αIIbβ3 structural studies that used “traditional” approaches to purify the protein with affinity chromatography, we believe our minimum enrichment, with only size-exclusive chromatography, is the key to preserving this unique homodimer confirmation. Further, the additional homodimer interface formed by two αIIbβ3 molecules stabilizes the protein in its intermediate state before the full extension. Our working hypothesis is that homodimerization is a previously unknown self-regulatory mechanism to temporally regulate its ligand accessibility in the intermediate state. (**Figure 7**) Future studies are needed to validate this conformation in live cells and to determine its impact on αIIbβ3 function. However, we do currently have indirect evidence to support our hypothesis. Both extended αIIbβ3 molecules in the intermediate conformation and in the homodimer conformation were unliganded, which is distinct from the bent state. (**Supplemental Figure 4**) Additionally, this homodimer conformation of αIIbβ3 may open a new avenue for antiplatelet therapy, suggesting a potential focus on the homodimer interface rather than directly targeting the ligand binding site of αIIbβ3.

Our results highlight the potential of cryo-EM coupled with BaR for system structural proteomics. BaR is an unbiased and untargeted approach. This method could be a double-sided sword that not only brings opportunities to simultaneously solve novel structures of proteins in different states from crude samples but also brings in inherent limitations. First, it is a challenge to solve the structure of a particular protein with low abundance in the crude sample. Second, the nature of the method makes it difficult to identify proteins with electron density maps lower than 4 Å resolution. Advances in cryo-EM sample preparation and the BaR methodology will overcome these limitations in future studies.

In conclusion, our study was the first to focus on platelets using a structural-omics approach that combines cryo-EM with Build-and-Retrieve data analysis. Using this approach, we were able to demonstrate the highly structural dynamics of αIIbβ3 from native platelet membranes, which actively rearrange among three different states: bent, intermediate, and homodimer. In addition, we showed the strength of the BaR methodology in preserving natural regulatory cues of the proteins from their environment. Our study highlights the potential of creating a structural atlas of platelet membrane protein complexes, which will ultimately enrich our understanding of the foundation of platelet signaling circuitry in response to surrounding, epigenetic, and genetic changes.

## Acknowledgements

MN receives research funding from the National Institutes of Health (NIH) and the National Heart Lung and Blood Institute (NHLBI) (HL098217 and HL154026). XH receives funding from the American Heart Association (AHA) Postdoctoral Fellowships (897185) and the American Society of Hematology (ASH) Fellow Scholar Award. This research was also supported in part by the grant AI145069 to EY from the National Institute of Allergy and Infectious Diseases (NIAID). We thank Belinda Willard and Ling Li at Lerner Research Institute’s Proteomics & Metabolomics Core for the acquisition of mass spectrometry data. The timsTOF Pro2 mass spectrometer instrument was purchased via an NIH-shared instrument grant, S10 OD030398. We are grateful to the Cryo-Electron Microscopy Core at the CWRU School of Medicine and Drs. Kunpeng Li and Kyle Whiddon for access to the sample preparation and Cryo-EM instrumentation.

## Data availability

Coordinates and EM maps for three conformations of integrin αIIbβ3 can be found at PDB accession numbers 9E8A, 9E8B, 9E8C and EMDB accession codes EMD-40713, EMD-40715, EMD-40716, EMD-40686, EMD-40688, and EMD-40691, respectively. The MS proteomics data have been deposited to the ProteomeXchange Consortium (http://proteomecentral.proteomexchange.org) *via* the PRIDE partner repository with the data set identifier PXD058005.

## Author Contribution

XH, EY, and MN conceived the study. XH and MN designed the experiments. XH, CS, ZZ, ML, MM, and MN performed the experiments and analyzed the data. XH and MN wrote the manuscript. CS, ZZ, ML, MM, and EY critically read and edited the manuscript.

---

**Movie 1.** Local motion and dynamics of the flexible αIIbβ3 were demonstrated by 3DFlex function embedded in the CryoSparc v4.6.0 suite.

## Supplemental Materials and methods

### Preparation of human platelets

Blood was collected from healthy donors using butterfly needles into sodium citrate. Platelet-rich plasma (PRP) was obtained by centrifuging at 200g for 10 minutes followed by 10-minute incubation, both at room temperature. Platelet concentrations were determined using a Coulter Counter. Platelets were collected by an additional centrifugation at 2200g for 16 minutes.

### Cryo-EM sample preparation

The pelleted human platelets were directly lysed by sonication in lysis buffer (phosphate buffer, 5 mM, pH 7.4) with protease inhibitor (Roche). The membrane fraction was collected by ultracentrifugation at 185,000g for 60 minutes at 4 °C. The membrane fraction was washed with high salt buffer (20 mM HEPES pH 7.5, 1M NaCl, 20 mM KCl, 10 mM MgCl_2_) followed by low salt buffer (20 mM HEPES pH 7.5, 50 mM NaCl). The washed platelet membrane fraction was homogenized and solubilized in solubilization buffer (1% DDM, 0.1% CHS, 20 mM HEPES pH 7.5, 50 mM NaCl, 10% glycerol) with protease inhibitor cocktail overnight at 4 °C. The insoluble fraction was removed by ultracentrifugation at 185,000g for 60 minutes at 4 °C. Sample was then concentrated, passed through a 0.22 μm centrifugal filter, and enriched using size-exclusion chromatography (Superose 6 increase 10/300, GE Healthcare). Protein sizes corresponding to 200–550 kDa were collected and assembled into lipidic nanodiscs (MSP1E3D1; POPC:POPZ:POPG). The sample was then passed through a 0.22 μm centrifugal filter and enriched using size-exclusion chromatography (Superose 6 increase, GE Healthcare). Protein sizes corresponding to 200–350 kDa were used for cryo-EM analysis. A sample of 2.5 μL (5 mg/mL) was applied to Quantifoil R 1.2/1.3 Cu 300 holey grids. Samples were blotted for 16 s and plunged frozen into liquid ethane using a Vitrobot (Thermo Fisher). The resulting grids were stored in liquid nitrogen until data collection.

### Data collection

A Titan Krios equipped with a K3 direct electron detector was used to collect data in super-resolution mode at 81,000 magnifications (physical pixel size of 1.070 Å/pix, 0.535 Å /pix super-resolution). Data were collected in correlated-double sampling mode using Serial EM.

### Data processing

Data were processed in the cryoSPARC V4.6.0 suite using the BaR protocol as described previously.^1–5^ The detailed data processing strategies are described in the Supplemental Data.

### Protein Identification, model building and refinement

The near-atomic-resolution cryo-EM maps were used for protein identification using the online server DeepTracer^TM^ and the program phenix.sequence_from_map in the PHENIX suite.^3,6,7^ Initial models were built using the AlphaFold ^8,9^ via the UniProt database and aligned to the cryo-EM maps in Chimera. Models were refined using phenix.real_space_refine and Coot. Final structures were evaluated using MolProbity. 3D FSC was used to quantitatively evaluate each cryo-EM map at distinct viewing directions.

### Quantification and statistical analysis

Standard GS-FSC (Gold Standard-Fourier Shell Correlation) curves at a threshold of 0.143 were computed using cryoSPARC v4.6.0 to obtain the final resolutions of protein models. The final atomic models were evaluated using MolProbity.

### Proteomic analysis of platelet membrane fractions

The platelet membrane proteins in lipidic nanodiscs from the size-exclusion chromatography enrichment was digested by trypsin. The resulting peptides were desalted by a C18 Microspin column (Nest Group, Ipswich, MA) per the manufacturer’s instructions and analyzed by LC-MS/MS using a ThermoScientific Fusion Lumos mass spectrometry system. Proteins were identified by comparing all experimental peptide MS/MS spectra against the UniProt human database using the Andromeda search engine integrated into the MaxQuant version 1.6.3.3. Carbamidomethylation of cysteine was set as a fixed modification, whereas variable modifications included oxidation of methionine to methionine sulfoxide and acetylation of N-terminal amino groups. For peptide/protein identification, strict trypsin specificity was applied, the minimum peptide length was set to 7, the maximum missed cleavage was set to 2, and the cutoff false discovery rate was set to 0.01. Match between runs (match time window: 0.7 min; alignment time window: 20 min) and label-free quantitation (LFQ) options were enabled. The LFQ minimum ratio count was set to 2. The remaining parameters were kept as default.

---

**Supplemental Figure 1.**
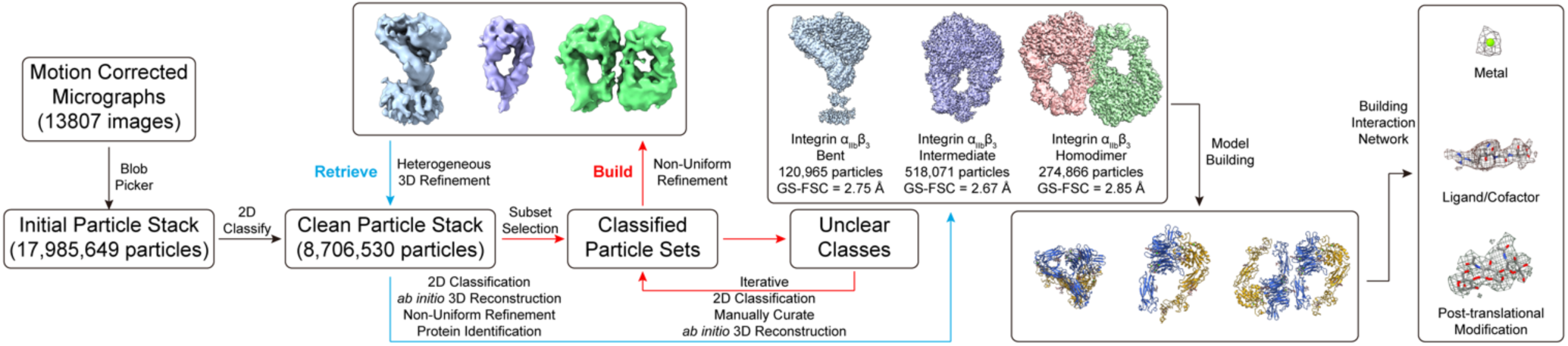
The Build-and-Retrieve (BaR) data processing method workflow. An unbiased blob picker was applied on 13807 motion-corrected micrographs to generate 17,985,649 initial particles. The 2D classification was used to generate a subset of a clean particle stack that contains 8,706,530 particles. Classes of particles that contain clear 2D features were used to *Build* the low-resolution initial models, which were used to *Retrieve* more particles from the clean particle stack. The improved maps at ∼3 Å resolution were used for protein identification. Final maps were used for model building and refinement.

**Supplemental Figure 2.**
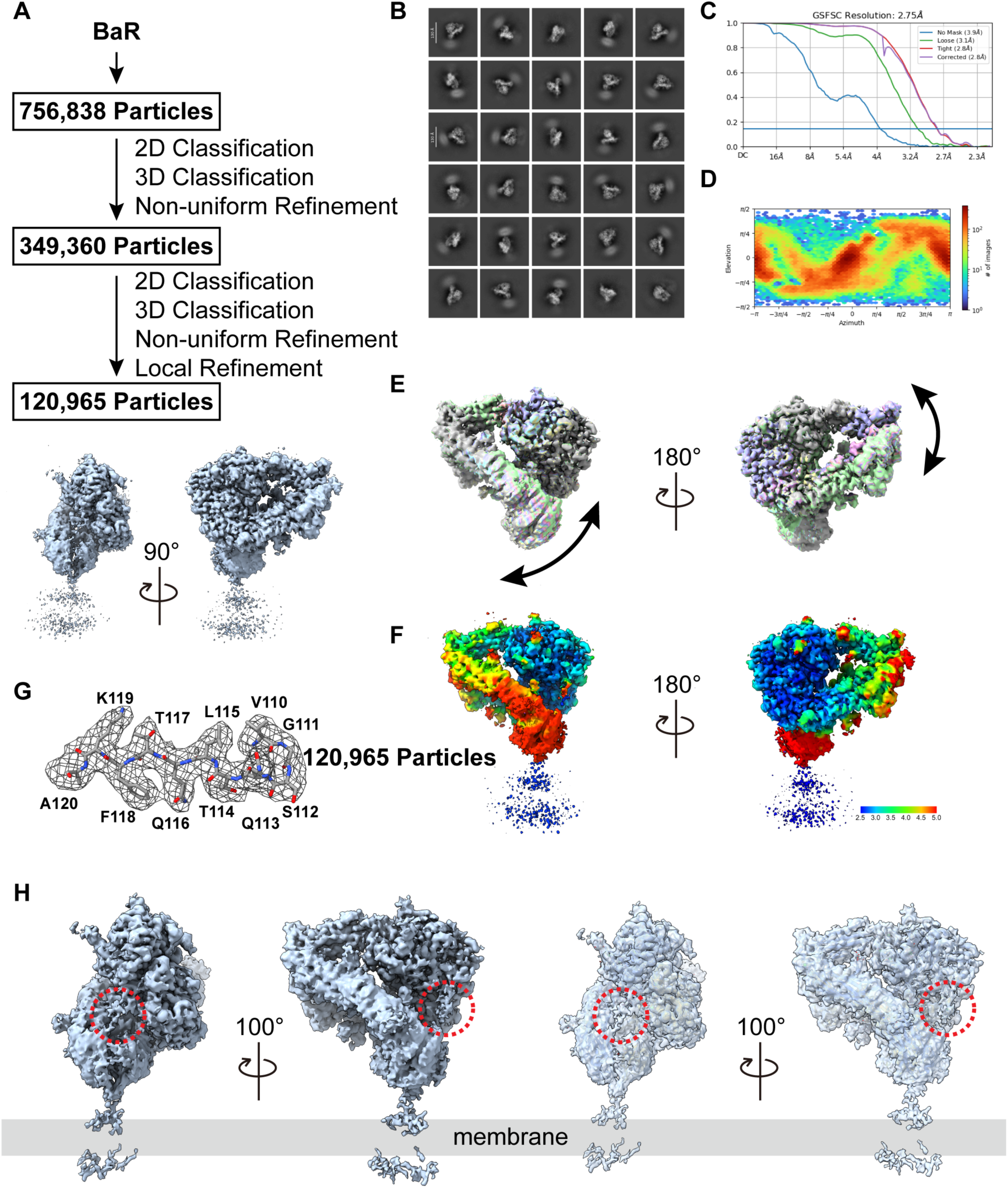
Cryo-EM analysis of bent integrin αIIbβ3. **A.** Bent integrin αIIbβ3 particles processing flowchart. **B.** Representative 2D classes of bent integrin αIIbβ3. **C.** Fourier Shell Correlation (FSC) curves. **D.** Angular distribution calculated from cryoSPARC for particle projection. **E.** 3DFlex analysis shows the domain dynamic of integrin αIIbβ3 in the bent conformation. **F.** Cryo-EM density map of bent integrin αIIbβ3 colored by local resolution. **G.** Local density map of bent integrin αIIbβ3. **H.** Cryo-EM density map of bent integrin αIIbβ3 at a contour level of 0.08 was shown on the left two panels. The same map with the model overlay was shown on the two right panels. The red circle highlighted the extra ligand density extended from the binding pocket that contains the RGD motif.

**Supplemental Figure 3.**
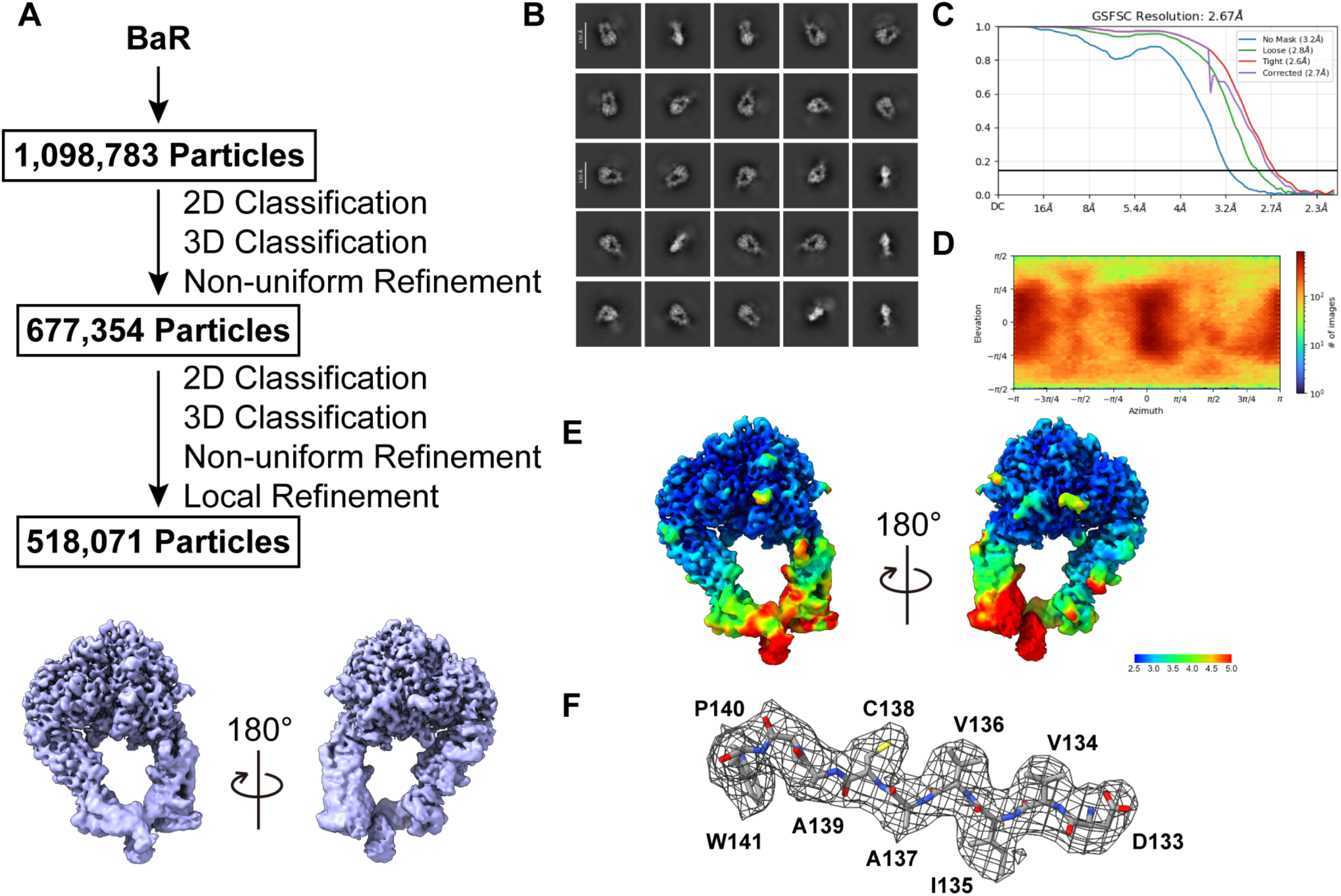
Cryo-EM analysis of intermediate integrin αIIbβ3. **A.** Intermediate integrin αIIbβ3 particles processing flowchart. **B.** Representative 2D classes of intermediate integrin αIIbβ3. **C.** Fourier Shell Correlation (FSC) curves. **D.** Angular distribution calculated from cryoSPARC for particle projection. **E.** Cryo-EM density map of intermediate integrin αIIbβ3 colored by local resolution. **F.** Local density map of intermediate integrin αIIbβ3.

**Supplemental Figure 4.**
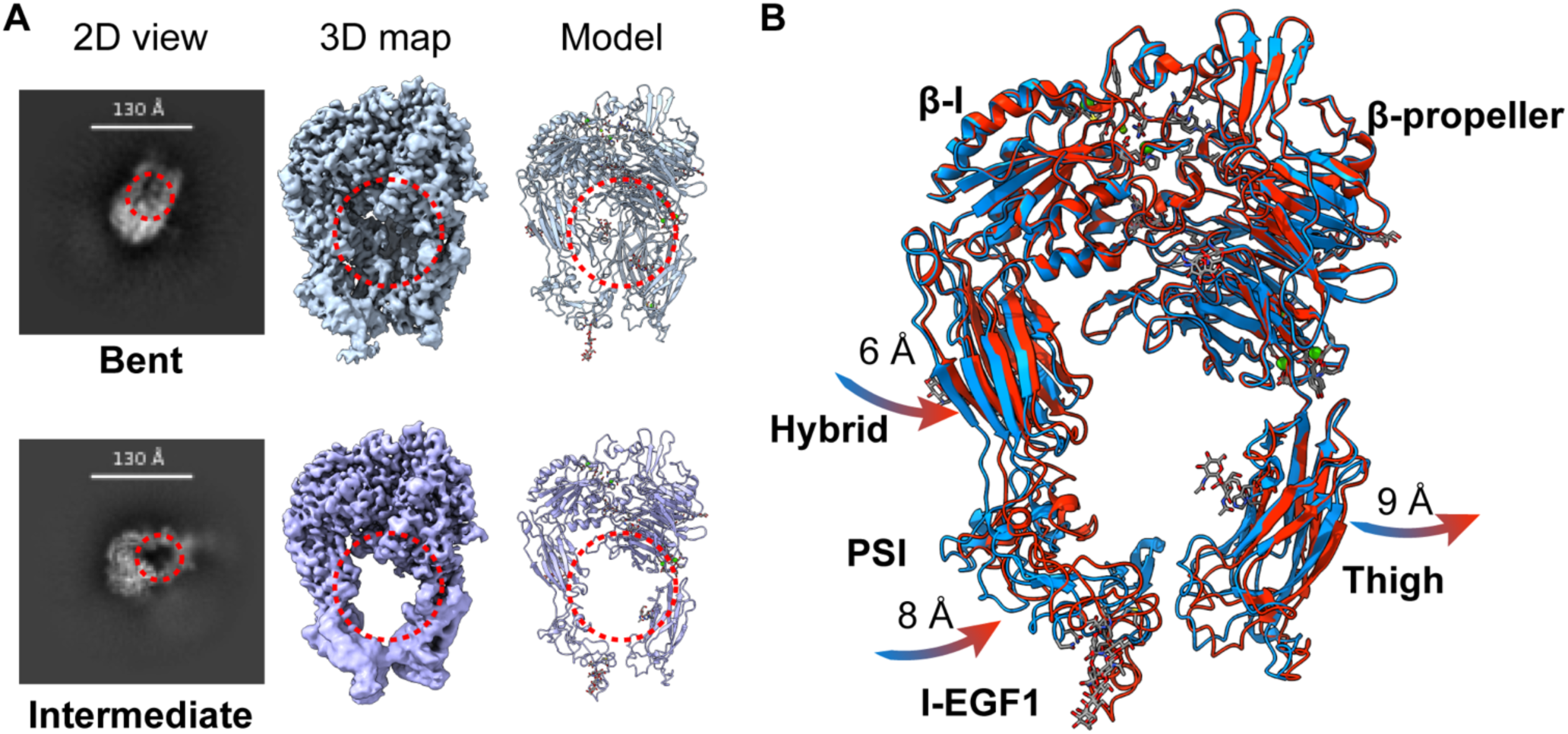
Comparison of the integrin αIIbβ3 in bent and intermediate conformations. **A.** An additional density (red circle) can be observed in the 2D projection of the top view of the bent conformation but not in the intermediate state. This is the density of the lower leg region of the molecule, seeing through the headpiece in the 3D map and the final model. **B.** Overlay of the bent (blue) and intermediate (red) headpieces conformations demonstrates the domain movement between states.

**Supplemental Figure 5.**
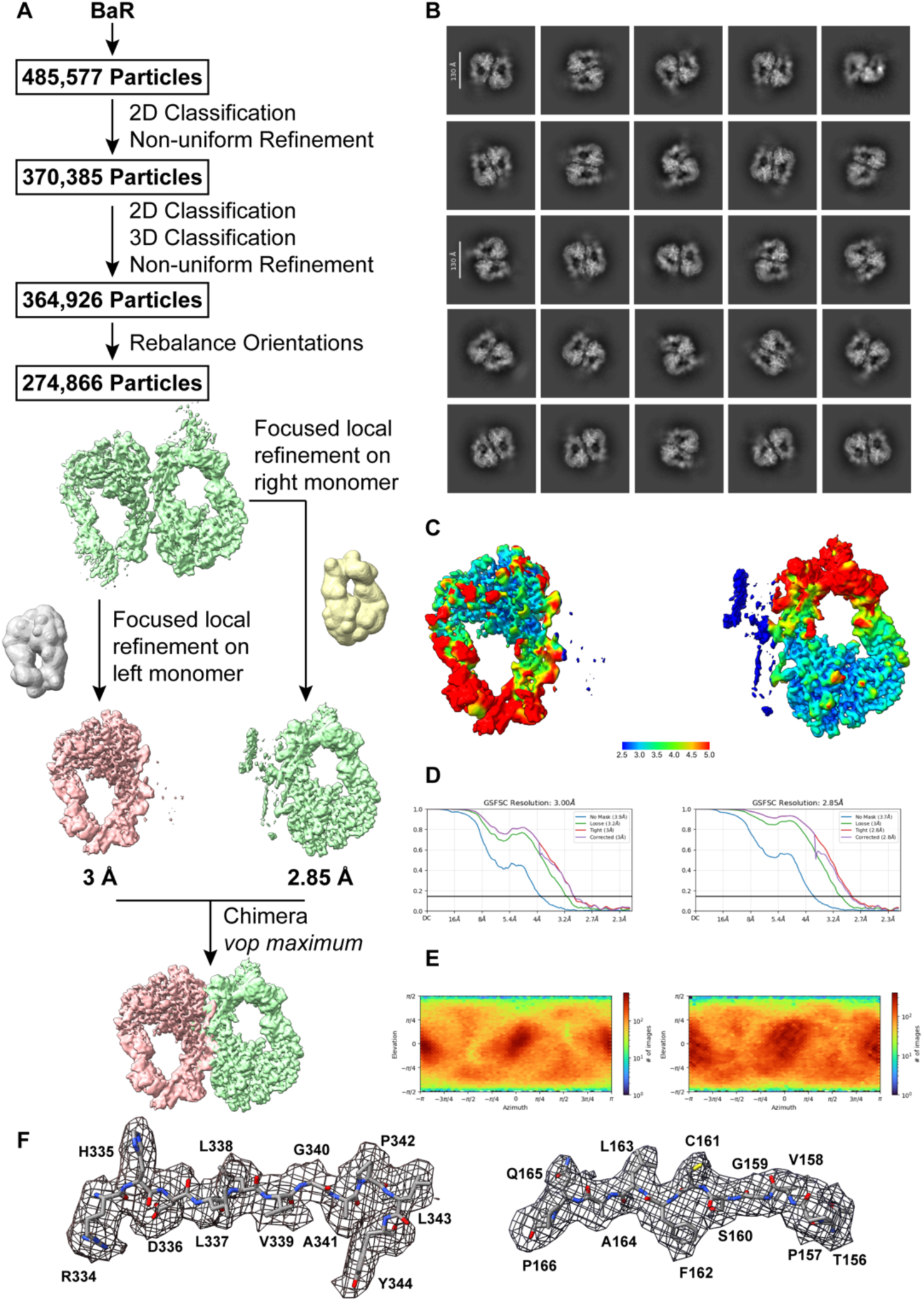
Cryo-EM analysis of integrin αIIbβ3 homodimer. **A.** integrin αIIbβ3 homodimer particles processing flowchart. **B.** Representative 2D classes of dimer integrin αIIbβ3. **C.** Cryo-EM density map of dimer integrin αIIbβ3 colored by local resolution. **D.** Fourier Shell Correlation (FSC) curves. **E.** Angular distribution calculated from cryoSPARC for particle projection. **F.** Local density map of integrin αIIbβ3 homodimer.

